# An oral vaccine for SARS-CoV-2 RBD mRNA-bovine milk-derived exosomes induces a neutralizing antibody response *in vivo*

**DOI:** 10.1101/2022.12.19.517879

**Authors:** Quan Zhang, Miao Wang, Chunle Han, Zhijun Wen, Xiaozhu Meng, Dongli Qi, Na Wang, Huanqing Du, Jianhong Wang, Lu Lu, Xiaohu Ge

**Affiliations:** Tingo Exosomes Technology Co., Ltd, Tianjin, China; Tingo Regenerative Medicine Technology Co., Ltd, Tianjin, China

**Keywords:** bovine milk-derived exosomes, SARS-CoV-2, receptor binding domain, mRNA, oral vaccines, neutralizing antibody

## Abstract

The severe acute respiratory syndrome coronavirus type 2 (SARS-CoV-2) that causes the coronavirus disease 2019 (COVID-19) has presented numerous challenges to global health. The vaccines, including lipid-based nanoparticle mRNA, inactivated virus and recombined protein, have been used to prevent SARS-CoV-2 infections in clinics and are immensely helpful against the epidemic. Here, we first present an oral mRNA vaccine based on bovine milk-derived exosomes (milk-exos), which encodes the SARS-CoV-2 receptor binding domain (RBD) as an immunogen. The results indicated that RBD mRNA delivered by milk-derived exosomes can produce secreted RBD peptide in 293 cells *in vitro* and stimulated neutralizing antibodies against RBD in mice. These results indicated that bovine milk-derived exosome-based mRNA vaccine could serve as a new strategy for preventing SARS-CoV-2 infection. Meanwhile, it also can work as a new oral delivery system for mRNA.

**One Sentence Summary:** Oral SARS-CoV-2 mRNA vaccine based on bovine milk-derived exosomes can stimulate neutralizing antibodies in mice.

## Introduction

COVID-19 typically presents symptoms common to many respiratory infections, including fever and cough. Still, in many cases, it progresses to a more severe disease that may include acute respiratory distress, disseminated disease, and death (1–3). The SARS-CoV-2 Spike protein is a homotrimeric transmembrane glycoprotein composed of S1 and S2 subunits. When spike protein interacts with specific receptors on host cells, its receptor binding domain (RBD) is located at the C-terminus of S1 and can specifically bind to the receptor angiotensin-converting enzyme 2 (ACE2) and initiates the membrane fusion between the virus and host cell (4–7). Studies have shown that RBD is the main target of most of the neutralizing activity in immune serum, suggesting that RBD may be a potential target for the 2019-nCoV vaccine or therapy (8–10).

Currently, the available platforms for COVID-19 vaccines include inactivated vaccines, live attenuated vaccines, recombinant protein vaccines, viral vector vaccines, and nucleic acid vaccines (11). In 2021, the revenue of COVID-19 vaccine products of global key enterprises showed that Pfizer and German BioNTechnology’s mRNA vaccine revenues were 367.8 billion dollars and 213.6 billion dollars, respectively, accounting for more than half of the global vaccine market share. Pfizer’s mRNA vaccine accounted for the highest proportion because it rapidly attracted great attention and promotion for its safety, high efficiency, and short production cycles. Compared with other vaccine technology platforms, the mRNA vaccine platform also has these advantages, including 1. an excellent immune effect; 2. a short R&D cycle; 3. convenient large-scale production (12, 13); 4. no risk of infection or genome integration (14–16); 5. the human expression system expresses antigens with good immunogenic recovery (17); 6 the stimulation of humoral and cellular immunity (18, 19); 7. no adjuvant, and so on. However, mRNA vaccines are also flawed, with high technical barriers and delivery technology patent restrictions; they are unstable and inconvenient to preserve and transport. Traditionally, it needs an intramuscular injection and a professional operation.

Most vaccines now undergoing clinical trials are injected intramuscularly or subcutaneously, restricting immune activation to the few draining lymph nodes (20, 21). Oral administration is thought to have a higher safety profile, better patient compliance, and lower medical costs than injection (20, 22). The difficulties of the complex gastrointestinal environment and intestinal epithelial barriers have limited the use of oral vaccines. To interact with the abundant immune cells in the lamina prima, an ideal oral vaccine must tolerate the gastrointestinal environment and overcome intestinal epithelial barriers (23).

Exosomes are cell-derived, membranous vesicles present in nearly all bodily fluids. With sizes ranging between 35-120 nm in diameter, exosomes are composed of a phospholipid bilayer derived from the membrane of the cell of origin. It has recently gained attention due to its natural role of shuttling molecular cargos (e.g., DNA, small RNAs, proteins, and lipids) between distant cells in the body. It has been confirmed that bovine milk is rich in exosomes and exhibits the similar potential to serve as drug-delivery nanocarriers (24). Milk is a more affordable and accessible source compared to cell culture media. Moreover, milk-exos may provide additional benefits as naturally desirable oral delivery carriers, which indicates that milk-exos constitute a more convenient and patient-friendly therapeutic modality (25). Taking into account the biological properties of milk exosomes and overcoming the technological challenges of mRNA vaccines, we have currently described a method for creating milk exosomes that are loaded with RBD mRNA, evaluated their effectiveness in delivering functional mRNA, developed a novel oral vaccine technology platform based on milk exosomes and investigated the preliminary efficacy of the novel oral vaccine based on milk exosomes in mice to elicit humoral immunity to SARS-CoV-2 spike proteins.

## Results

### Preparation of bovine milk-derived exosomes

To evaluate our preparation methods, bovine milk-derived exosomes (milk-exos) were isolated and purified by density gradient ultracentrifugation (DC) (Fig. 1A). Six components were collected by DC, among which F1 was about 5 mL, and F2 to F6 were all 7 mL (Fig. 1B). To characterize the isolated milk-exos biophysically, biomarkers (CD9, TGS110), and morphology was used to determine the exosomes containing fraction. Exosomes were mainly concentrated in F3 and F4 (Fig. 1C, D). The morphology exhibited a rich profusion of mixed populations of exosomes with predominantly intact vesicles consistent with classical exosome-like morphology and a typical cup-like structure (Fig. 1D). Milk exosomes are characteristic of exosomes in the range of 30-150 nm in diameter were observed in 0.95 mol/L-1.30 mol/L sucrose. As fractions 3 and 4 were enriched in exosomes, they were pooled together for further analysis.

**Figure 1.**
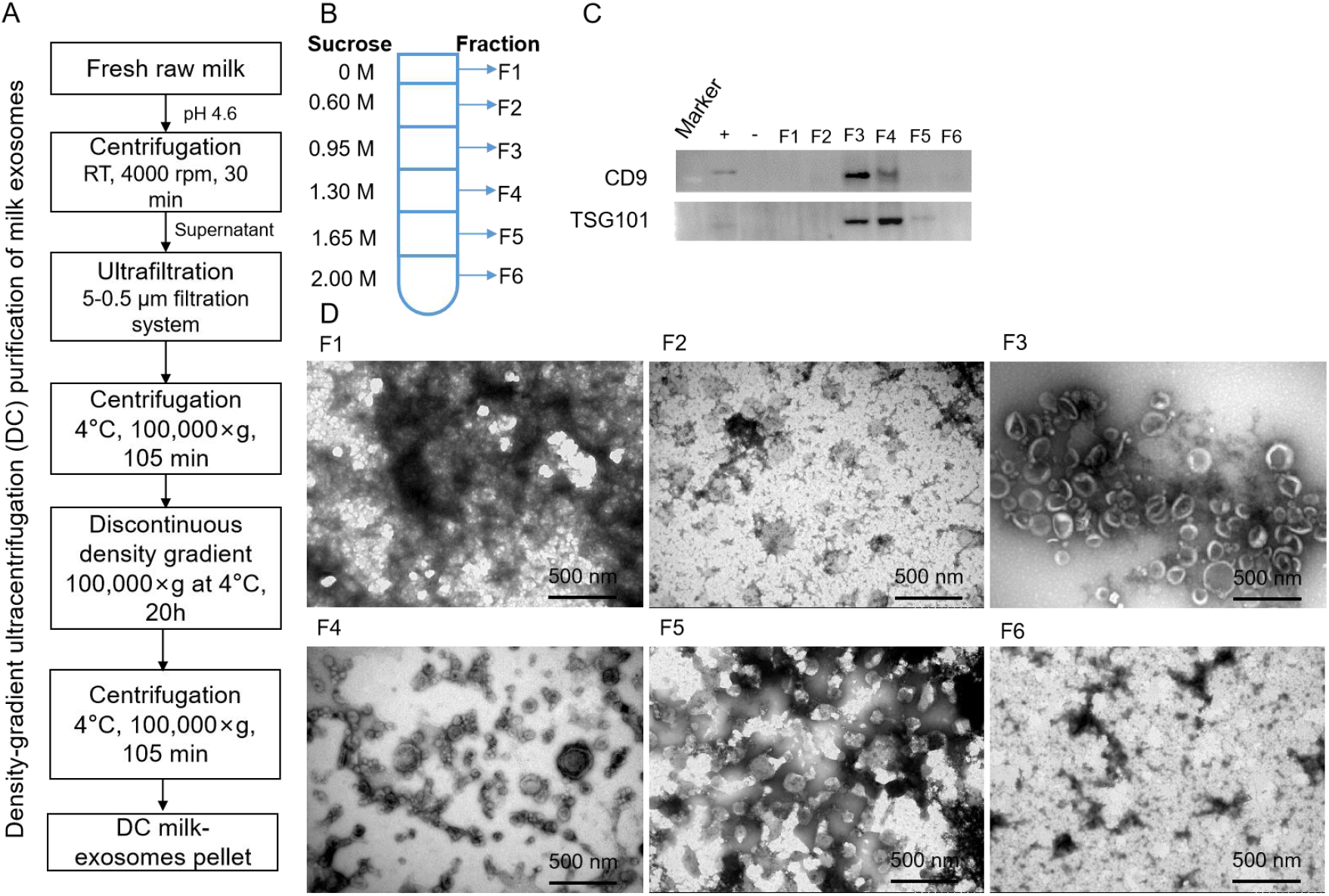
Identification of bovine milk-derived exosomes in fractions of density gradient ultracentrifugation. (A) Schematic representation of the major steps involved in isolating of exosomes from bovine raw milk. (B) F1–F6 represents the corresponding concentrations of sucrose. (C) The exosome suspension was analyzed by western blot. Immunoblots showed exosomes marker in different milk exosome fractions. +, positive control (the protein of HaCat cells); -, negative control (the protein of Hela cells). (D) The morphology of different fractions obtained by TEM. (Scale bars = 500 nm).

### Purification and characterization of bovine milk-derived exosomes by density gradient ultracentrifugation

The morphology of pooled exosomes fraction exhibited a rich profusion of exosomes consistent with classical exosome-like morphology (Fig. 2A), size distribution (Fig. 2B) and protein markers (Figure 2C). A three-way Venn diagram of proteins revealed 1022 proteins common to all datasets, and 961 proteins were commonly identified in all three DC-milk-exos, as shown in Figure 2C. Proteomics analysis of the samples showed that exosome protein lysates were prepared and verified with cluster analysis for vital exosomal membrane markers CD9, CD63, CD81, and TSG101. The absence of the microvesicle surface markers GM130 and calnexin, as well as the absence of the endoplasmic reticulum (ER) marker calnexin, confirmed that the isolated milk-exos were not contaminated with other multivesicular bodies (Fig. 2C). Three batches of bovine milk-derived exosomes obtained by density gradient ultracentrifugation (DC-milk-exos) were analyzed and indicated that there were no differences between multiple batches of DC-milk-exos.

**Figure 2.**
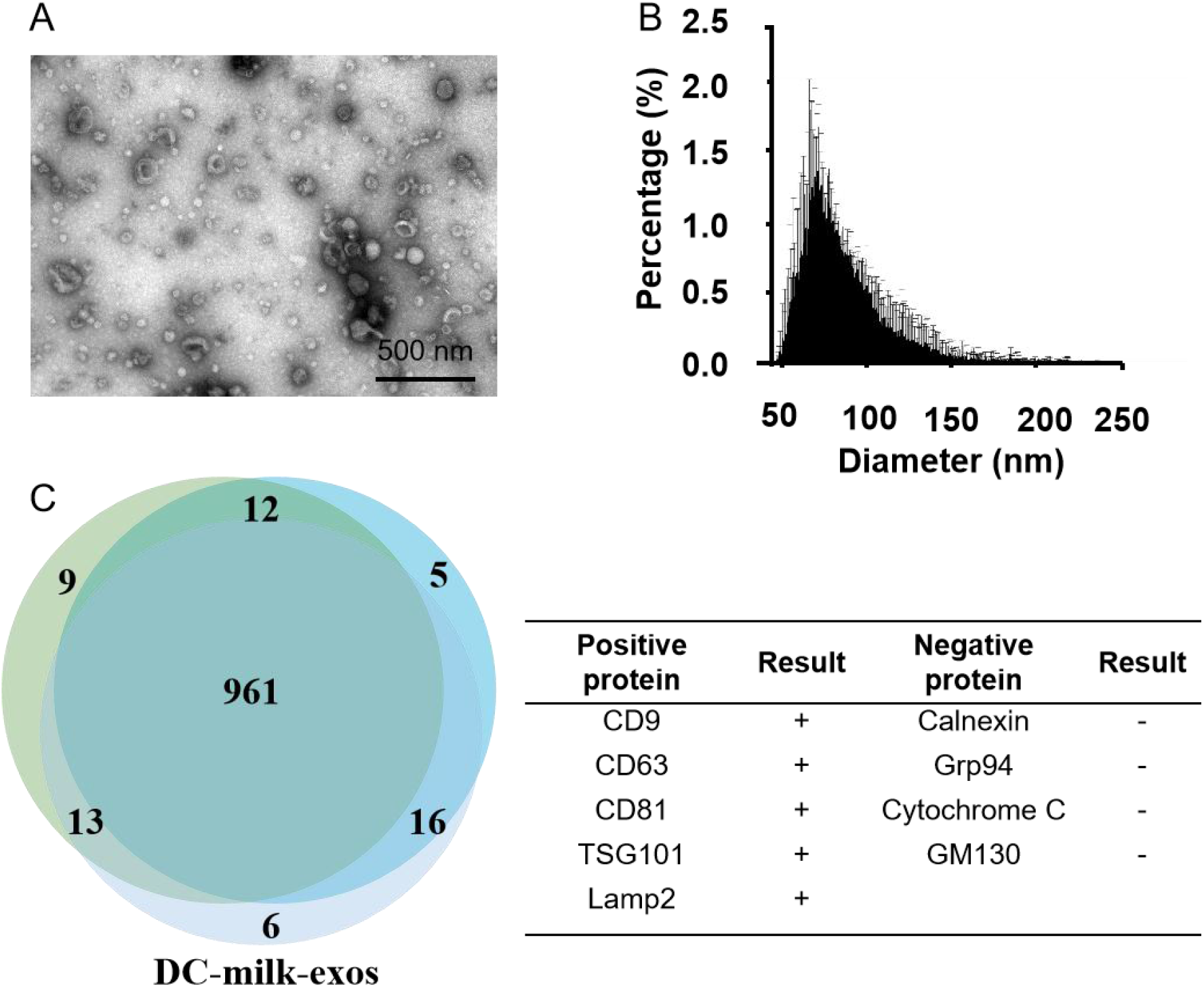
Characterization of DC-milk-exos. (A) The morphology and (B) the particle size analysis were detected by TEM and nanoFCM, respectively. (C) A three-way Venn diagram of proteins from three batches of DC-milk-exos revealed 1022 proteins common to all datasets. Cluster analysis for vital exosomal membrane markers CD9, CD63, CD81, TSG101, microvesicle surface markers GM130, and endoplasmic reticulum (ER) marker calnexin are indicated in the table. Abbreviations: TEM, transmission electron microscope; DC-milk-exos, bovine milk-derived exosomes by density gradient ultracentrifugation. Scale bars = 500 nm.

### Loading of RBD mRNA Milk derived exosomes

To determine whether DC-milk-exos could be loaded with exogenous, *in vitro* synthesized mRNAs, we designed and synthesized a test receptor binding domain (RBD) mRNA encoding immunogenic forms of the SARS-CoV-2 spike. The RBD coding sequence region (CDS) is 675 base pairs (bp) long and has the FLAG tag. Examination of the *in vitro* synthesized RBD mRNA using a bioanalyzer (Agilent) confirmed that the RBD mRNA sample ran as a single band of 1100 bps (Fig. 3A), consistent with the size that we expected to design in vitro transcription. To assess its functionality, the RBD mRNA was transfected into 293T cells, and RBD peptide expression was interrogated the next day by western blot (Fig. 3B). These results indicated that RBD mRNA was synthesized according to in vitro transcription (IVT) and had a translational function that could be translated into RBD protein in cells.

**Figure 3.**
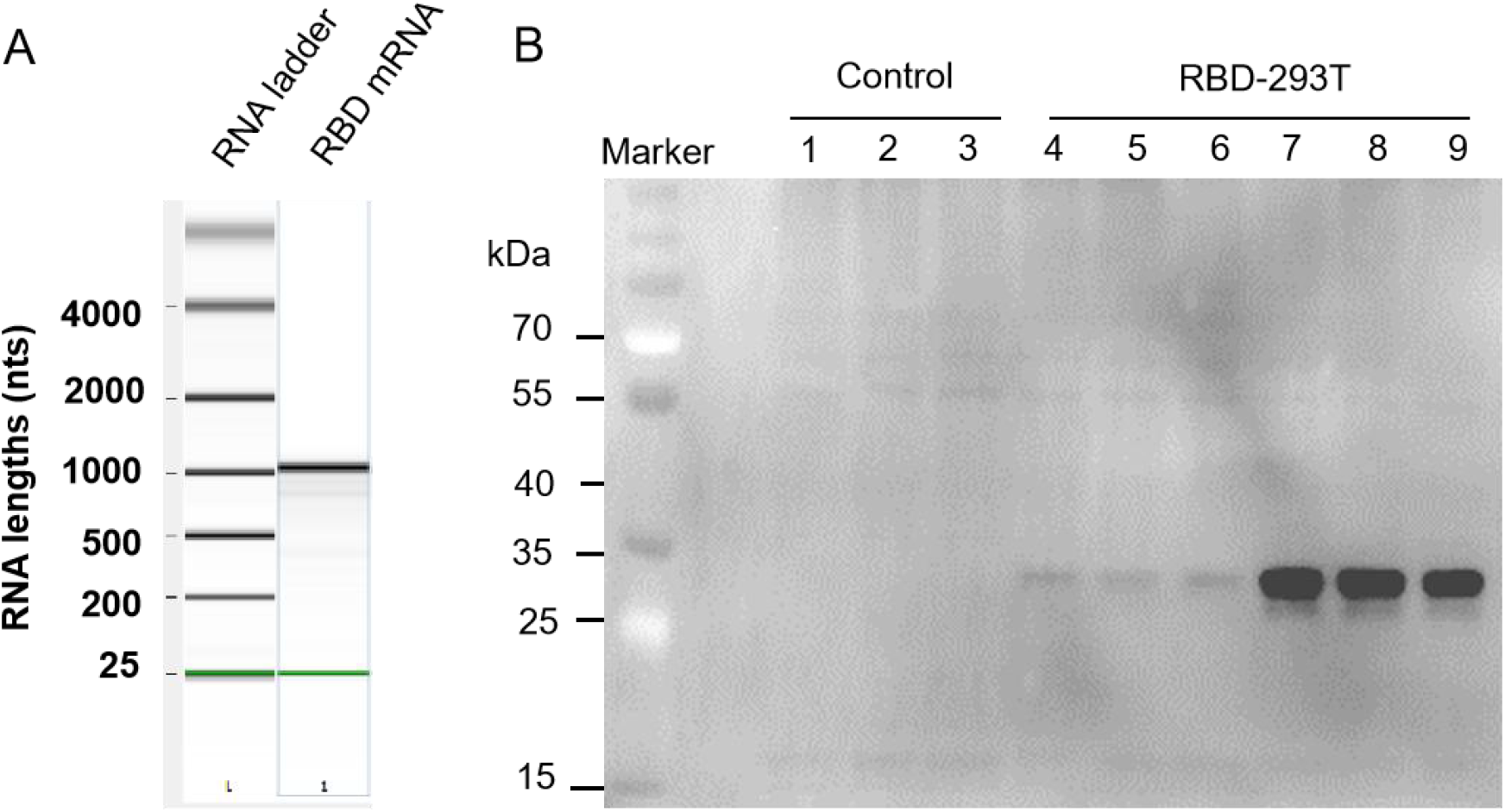
Characterization of SARS-CoV-2 receptor binding domain (RBD) mRNA. (A) Gel-like image of in vitro synthesized RBD mRNA interrogated using an RNA chip on an Agilent Bioanalyzer. Data for RNA markers, RBD mRNA, are presented from left to right. (B) Western blot for SARS-CoV-2 RBD protein expression in 293T cells when the RBD mRNA was transfected into 293T cells with an additional 24 h of treatment along with 1 μg and 3 μg RBD mRNA. Lanes 1-3: control; Lanes 4-6: 1 μg RBD-293T; Lanes 7-9: 3 μg RBD-293T.

To load the IVT RBD mRNA into DC milk-exos, we first mixed it with cationic lipids (DOTAP) to generate lipid mRNAs. Then we loaded the lipid-mRNA into DC-milk-exos by mixing-induced partitioning (Fig. 4A). Both processes are driven by the attractive force of the charge, resulting in the encapsulation of lipid-mRNAs into Milk-exos membranes. To characterize the RBD mRNA Milk-exos biophysically, the morphology and size distribution and zeta potential analysis, were performed (Fig. 4B-D). To determine the loading efficiency of this process, the products of three independent mRNA-loading reactions were examined. A TaqMan-based real-time quantitative PCR assay showed that loading efficiency reached 57.3% (Fig. 4E).

**Figure 4.**
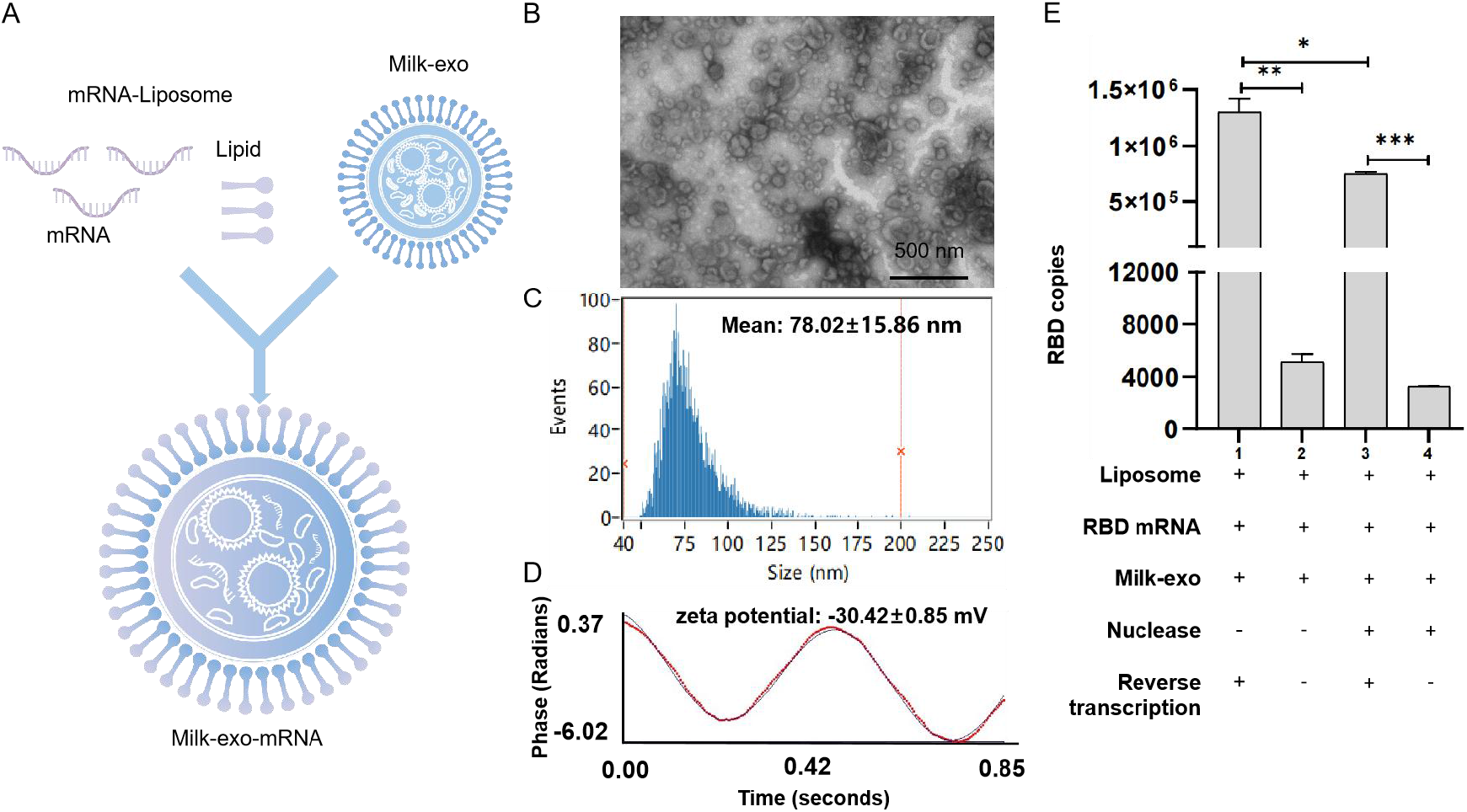
Characterization of milk-derived exosome-based vaccine for SARS-CoV-2. (A) Flowchart of vaccine preparation for SARS-CoV-2. The morphology (B), particle size distribution (C), zeta potential analysis (D), and RBD mRNA loading efficiency (E) were executed. Scale bars = 500 nm.

### Verification of oral vaccines for RBD mRNA-DC-milk-exos *in vitro* and *in vivo*

We next tested whether Milk-exos loaded with RBD mRNA could deliver functional RBD mRNA into human cells. Western blot and ELISA results established that RBD mRNA-loaded Milk-exos could deliver mRNA into 293 cells and produce RBD peptide 24 hr later. (Fig. 5A, B, p<0.01).

**Figure 5.**
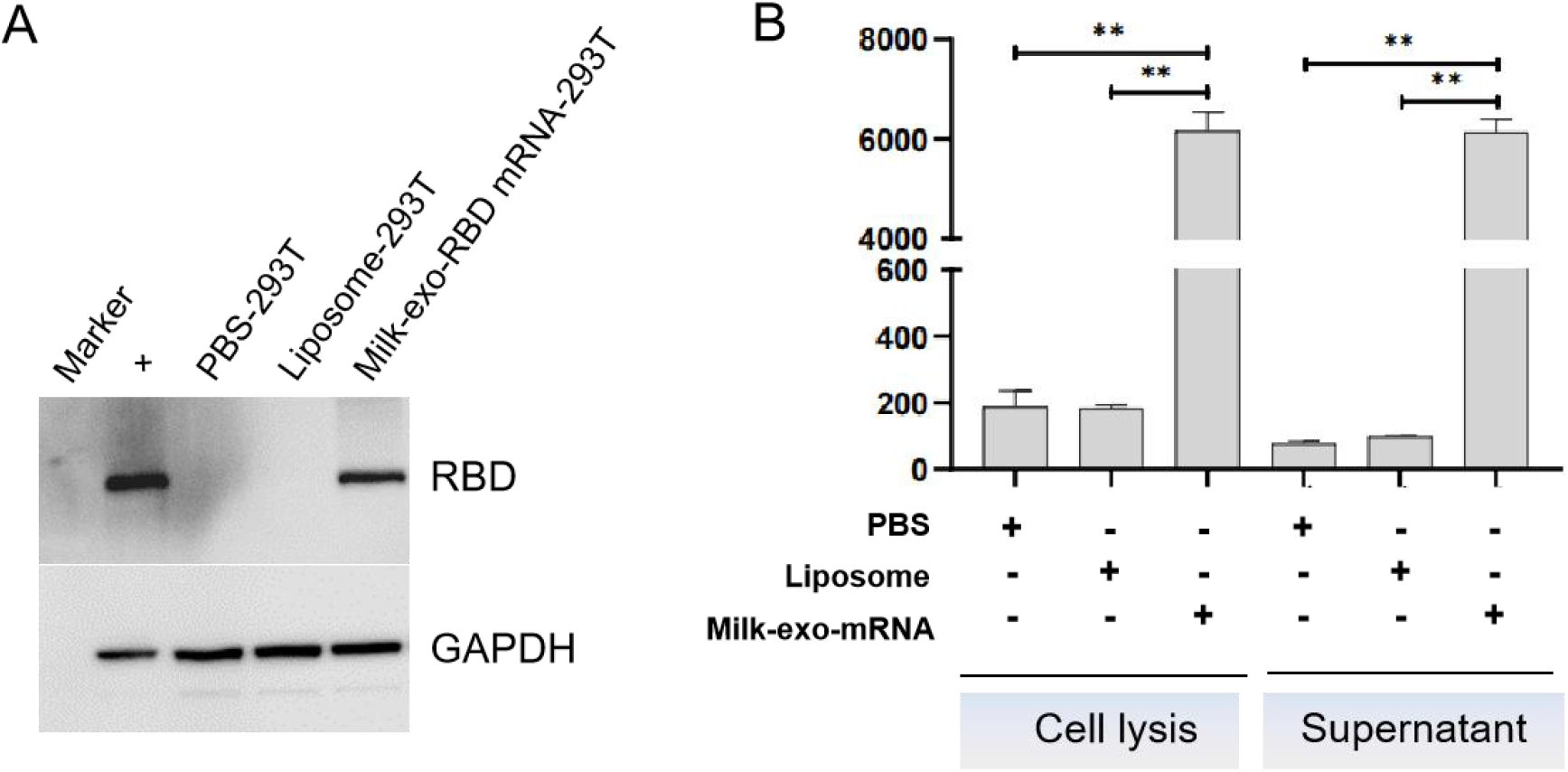
mRNA-loaded exosomes deliver functional SARS-CoV-2 RBD mRNA to human cells *in vitro*. (A) Western blot analysis of the expression of RBD mRNA delivered by oral vaccine in 293T cells. (B) The ELISA detection of RBD expressed in 293T cell lysate and secreted into the culture supernatant at 24 h after the milk-derived exosome-based vaccine transfection. The data are presented as mean standard deviation with a group size of three. \*\**P* < 0.01 vs. PBS.

To further confirm the ability to stimulate neutralizing antibodies, RBD mRNA-milk-exos were injected into the duodenum (i. d.) of 9-11-week-old female BALB/c mice (Fig. 6A). Blood (0.1 mL) was collected on days 0, 7, 14, 21, 28, 35, 42, and 49 for antibody detection before the animals were sacrificed. Using ELISA kits adapted for detecting mouse-derived antibodies, we observed that vaccinated animals produced a relatively constant level of neutralizing antibodies against RBD after the second injection (Fig. 6B, C).

**Figure 6.**
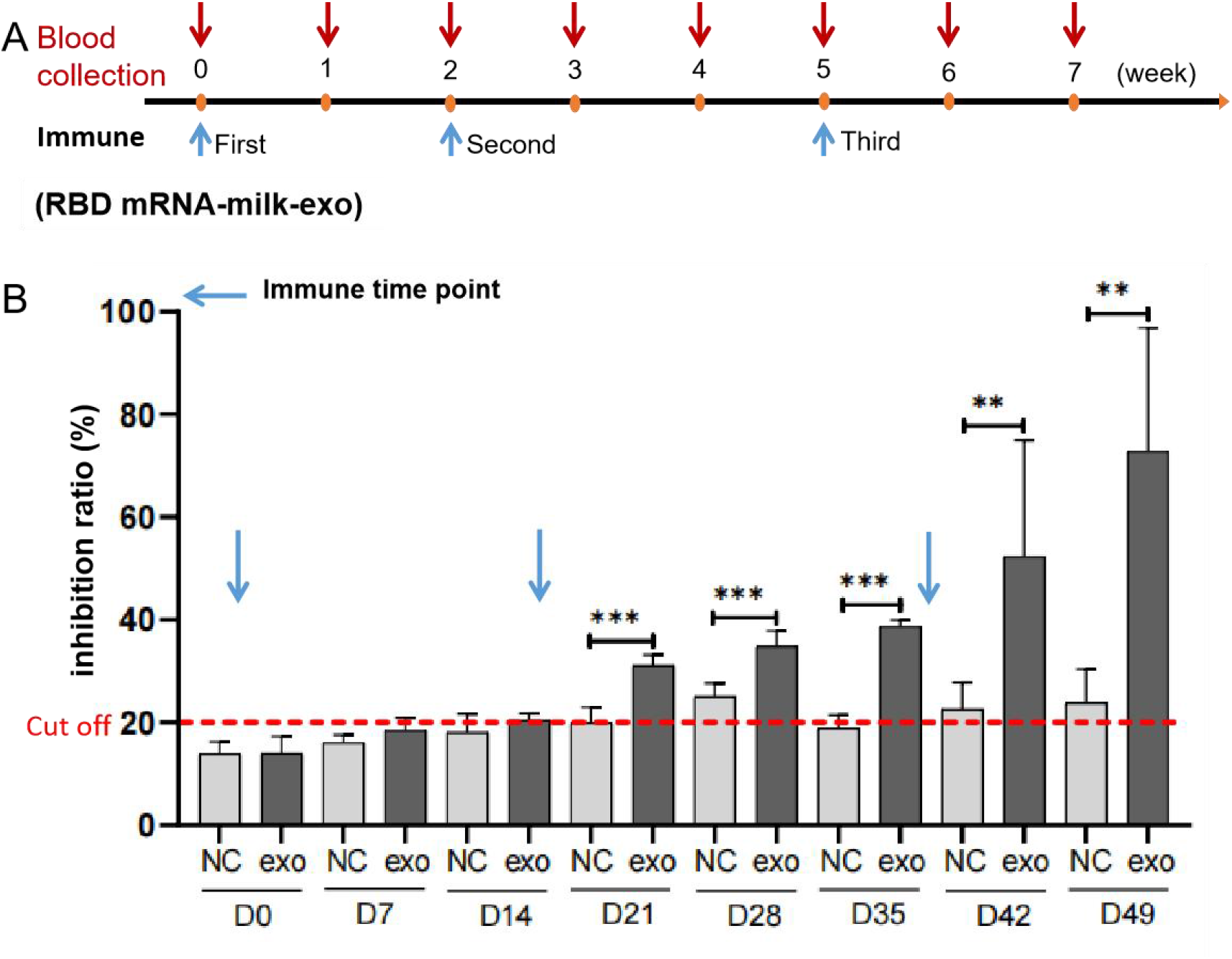
Validation of SARS-CoV-2 RBD mRNA-loaded milk exosomes delivering functional SARS-CoV-2 RBD mRNA *in vivo*. (A) Mouse immunization and sera sampling schedule. BALB/c mice received the same doses of an oral vaccine for the SARS-CoV-2-based milk-derived exosomes (N = 5) or the control saline (N = 3) on day 0 and were boosted again on days 14 and 35. Sera were collected on days 0 (pre-vaccination), 7, 14, 21, 28, 35, 42 and 49 (post-vaccination). The blue and red arrow represented the time points of immunization and blood collection, respectively. (B) The neutralizing antibodies of the oral vaccine for SARS-CoV-2-based milk-derived exosomes in serum were determined by ELISA. The red dotted line represented the cutoff value of neutralizing antibodies against the RBD peptide. N = 3 (control), N = 5 (RBD-DC-milk-exos), and the data are presented as mean±STD. \*\**P* < 0.01, \*\*\**P* < 0.001 vs. control group.

## Discussion

In this study, we demonstrated that an oral vaccine for SARS-CoV-2-based bovine milk-derived exosomes could deliver functional mRNA *in vitro* and *in vivo* and induce anti-S antibody responses. We first successfully verified the technical feasibility of oral mRNA delivery based on bovine milk-derived exosomes as delivery vehicles.

Compared with injection, oral administration was generally considered to have a better safety profile, better patient compliance, and lower medical costs (20, 22). Like liposomes, exosomes have a bi-lipid membrane and an aqueous core; therefore, they could potentially be loaded with hydrophilic and lipophilic drugs (26). However, the practical application of exosome-based therapeutics in clinical transformation remained an ongoing challenge. Many studies isolated exosomes from cell culture media with a low yield and cost, making scaling-up production difficult (27). A cost-effective and scalable source should be optimized based on the current limitations.

It was reported that bovine milk-derived exosomes exhibited a similar potential to serve as drug-delivery nanocarriers (24). Milk was a more affordable and accessible source compared to cell culture media. Moreover, bovine milk-derived exosomes might provide additional benefits as naturally desirable oral delivery carriers, which indicates that bovine milk-derived exosomes constitute a more convenient and patient-friendly therapeutic modality (25).

Herein, we presented an oral vaccine that differed from previous nanomaterials-based oral vaccine delivery technologies in that it comprised bovine milk-derived exosomes, and the milk-derived exosome-derived oral COVID-19 mRNA vaccine had its delivery technology and industrialization capability. Although the results from this study suggested the beneficial effects of an oral vaccine for the prevention of the novel coronavirus and offered a potential regimen for clinical application, we needed to expand the number of animals and further verify the efficacy of the oral vaccine in different animal models (such as large animals) in future studies. Furthermore, the precise molecular mechanisms and safety needed to be investigated further.

Although density-gradient ultracentrifugation has been the gold standard for exosome isolation and purification, there have been several studies on the extraction of exosomes from cow’s milk using differential centrifugation alone (28, 29) or precipitation (30, 31) techniques. In conjunction with differential centrifugation, we employed the density-gradient ultracentrifugation methods to isolate exosomes from milk that were essentially free of contamination from microvesicles. The final sample purity could reach 100%. In contrast, the technique used in this study to purify exosomes is incompatible with large-scale exosome production, which is one of the obstacles to the progress of industrialization. The good news is that we have established the chromatography-based novel manufacturing process to purify exosomes from bovine milk with better quality than production by DC (data are not shown).

It will come soon that an mRNA delivery system based on milk-derived exosomes will serve as a platform for mRNA therapeutics development in the recent future.

## Materials and methods

### Cell lines and cell culture

293T cells (SCSP-502) (human embryonic kidney epithelial cells) were purchased from the Cell Bank of the Typical Culture Preservation Committee, Chinese Academy of Sciences (Shanghai, China). 293T cells were grown in DMEM supplemented with 10% fetal bovine serum (FBS) and 1% penicillin/strep solution at 37 ° C and 5% CO2 concentration in a humidified atmosphere. The pooled cells were maintained in DMEM containing 10% (vol/vol) FBS, changing the medium every other day. When the confluence of cultured cells reached 80%, they were detached by treatment with 0.25% (wt/vol) trypsin and 0.1% (wt/vol) ethylenediaminetetraacetic acid (Gibco) and reseeded at a density of 1 × 10^4^ cells per cm^2^. Cultured cells before passage two were used for experiments. For RBD mRNA expression *in vitro*, cells were transfected with mRNA using Lipofectamine Messenger MAX, as suggested by the manufacturer (Thermo Fisher).

### Density-gradient ultracentrifugation purification of milk-exos (The DC-milk-exos)

Milk exosomes were isolated by density-gradient ultracentrifugation (DC). Briefly, the casein-free whey was centrifuged at 100,000 g (Beckman Coulter, USA) for 105 min to precipitate the milk-exos, resuspended in 1 mL PBS. The concentrated milk-exos were subjected to the top of a discontinuous density gradient consisting of 2 mol/L, 1.65 mol/L, 1.3 mol/L, 0.95 mol/L, and 0.6 mol/L sucrose (7 mL volume for each angle) in 250 mM Tris-HCl solution (pH 7.4). They were centrifuged at 100,000 g at 4 °C for 20h. The milk-exos fraction between fractions 3 (1.3 mol/L) and 4 (1.65 mol/L) was collected. To remove sucrose, the fraction was diluted in PBS to a final volume of 40 mL and centrifuged at 100,000 g at 4 ° C for 105 min. The pellet was resuspended in 1 mL of PBS. The DC-milk-exos were stored at −80 °C before use.

### Characterization of the DC-milk-exos morphology by transmission electron microscopy

The DC-milk-exos morphology was studied by transmission electron microscopy (TEM), as described before, with some modifications. First, the DC-milk-exos (100 μg/mL) were fixed by mixing with an equal volume of 4% (w/v) paraformaldehyde at room temperature for 15 minutes. The selected sample (10 μL) was then subjected to a formvar-carbon-coated TEM grid and kept at room temperature for 3 minutes. The grid was stained by adding 10 μL of uranyl oxalate solution (4% uranyl acetate, 0.0075 M oxalic acids, pH 7). The stained grid was investigated by a Hitachi 5600 plus transmission electron microscope operated at 120 KV and 50,000 magnification.

### Measurement of the particle size distribution of DC-milk-exos by NanoFCM

The particle size and number of milk-exos samples were characterized by the NanoFCM instrument (NanoFCM Inc., Xiamen, China) by following the operations manual. A silica nanosphere cocktail (Cat. S16M-Exo, NanoFCM Inc., Xiamen, China) containing a mixture of 68 nm, 91 nm, 113 nm, and 155 nm standard beads was used to adjust the instrument for particle size measurement. The instrumental parameters were set as follows: Laser, 10 mW, 488 nm; SS decay, 10%; sampling pressure, 1.0 kPa; sampling period, 100 μs; time to record, 1 min.

### Proteomics

The proteomics of DC-milk-exos was analyzed by LC–MS/MS using Easy NLC1200-Q Exactive and Fusion Lumos Orbitrap mass spectrometers (ThermoFisher), both fitted with nanoflow reversed-phase HPLC (Ultimate 3000 RSLC, Dionex). The nano-HPLC system was equipped with an Acclaim PepMap nano-trap column (Dionex-C18, 100 Å, 75 μm × 2 cm) and an Acclaim Pepmap RSLC analytical column (Dionex-C18, 100 Å, 75 μm × 25 cm). Typically, for each LC-MS/MS experiment, 1 μL of the peptide mix was loaded onto the enrichment (trap) column at an isocratic flow of 5 μL/min of 3% CH3CN containing 0.1% formic acid for 5 min before the enrichment column was switched in-line with the analytical column. The eluents used for the LC were 0.1% (v/v) formic acid (solvent A) and 100% CH3CN/0.1% (v/v) formic acid. The gradient used was 3% B to 25% B for 23 min, 25% B to 40% B in 2 min, 40% B to 85% B in 2 min, and maintained at 85% B for 2 min before equilibration for 10 min at 3% B before the next injection. All spectra were collected in positive mode using full-scan MS spectra scanning in the FT mode from m/z 300-1650 at resolutions of 70 000 (QE) and 120,000 (Lumos). A lock mass of 445.12003 m/z was used for both instruments. For MS/MS on the Lumos, the “top speed” acquisition mode (3 s cycle time) on the most intense precursor was used, whereby peptide ions with charge states ≥2 were isolated with an isolation window of 1.6 m/z and fragmented with HCD using a normalized collision energy of 35. For MSMS on the QE plus, the 15 most intense peptide ions with charge states ≥2 were isolated with an isolation window of 1.6 m/z and fragmented by HCD with a normalized collision energy of 35. A dynamic exclusion of 30 seconds was applied.

The raw files were searched using Proteome Discover (version 2.1, Thermo Fisher, Germany) with Sequest as the search engine. Fragment and peptide mass tolerances were set at 20 mDa and 10 ppm, respectively, allowing a maximum of 2 missed cleavage sites. The false discovery rates of proteins, peptides, and phosphosites were 1 percent. The differential expression proteins were analyzed by DAVID (Database for Annotation, Visualization, and Integrated Discovery) Bioinformatics Resource 2021 (http://david.abcc.ncifcrf.gov/) with recommended analytical parameters to identify the most significantly enriched signal transduction pathways in the data set.

### Western blot

Total cellular and milk-exos proteins were extracted with RIPA lysis buffer, and the protein concentration was determined by a BCA protein assay kit (Thermo, A53226). Following that, the protein was loaded onto SDS-polyacrylamide gel electrophoresis gels. After electrophoresis, proteins were transferred onto a polyvinylidene fluoride membrane and blocked in 5% (wt/vol) bovine serum albumin (BSA) for 1 hour at room temperature. Then the membrane was incubated with anti-TSG101 (Abcam, ab125011), anti-CD9 (Abcam, ab92726), anti-RBD (Abcam, ab277628), or anti-GAPDH (Proteintech, 60004-1) overnight at 4 °C. After washing in Tris-buffered saline with Tween (TBST), the horseradish peroxidase (HRP) secondary antibody was diluted 1: 10,000 with 5% (wt/vol) BSA and incubated with the membrane for 1 hour at room temperature. Excess secondary antibody was rinsed off the membrane with TBST, and a chemiluminescent signal was generated using the FluorChem E system (ProteinSimple, USA) according to the manufacturer’s protocol.

### Measurement of zeta potential

The Zeta potential of milk exosomes was measured thrice at 25 °C under the following settings: sensitivity of 85, a shutter value of 70, and a frame rate of 30 frames per second, while ZetaView software was used to collect and analyze the data.

### Construction of an oral vaccine for SARS-CoV-2 RBD-based DC-milk-derived exosomes (RBD-DC-milk-exos)

mRNA designed to express the RBD proteins was obtained from a commercial provider (Novoprotein). RBD mRNAs were purified using CIMmultus Oligo dT columns and resuspended in DNase- and RNase-free water using nuclease-free tips and tubes. Purified RBD mRNAs were prepared for loading into DC-milk-exos by preincubating them with cationic lipids, generating a lipid-mRNA product. Lipid-mRNAs were subsequently loaded into purified DC-milk-exos by mixing and incubation to construct an oral vaccine. The morphology, particle size distribution, and zeta potential of RBD mRNA-loaded DC-milk-exos were characterized by TEM, the NanoFCM instrument, and Zeta Pals, respectively. RBD mRNA loading efficiency was detected by Taqman-based quantitative real-time PCR (qRT-PCR). RBD mRNA expression was measured using western blot and enzyme-linked immunosorbent assay (ELISA).

### RNA extraction and Taqman-based RT-qPCR assay

Total RNA was extracted from the RBD-DC-milk-exos using TRIzol (Life Technologies, Carlsbad, CA). First-strand cDNA was synthesized using the PrimeScript^™^ RT Kit (TaKaRa, Beijing, China) and used as a template to determine the expression of RBD genes with the indicated primers and probes. The following primer sequences were used: RBD, forward 5’-CTCCAGGGCAA ACTGGAAAG-3’, and reverse 5’-AATTACCACCAACCTTAGAATCAAG-3’, probe, CCAGATGATTTTACAGGCTGCGTTATAG, using Premix Ex Taq™ (Probe qPCR) reagent (TaKaRa, Beijing, China) in qRT-PCR. The cycling parameters were as follows: a PCR reaction was carried out on 50 ng of cDNA samples using 0.2 μmol/L of each primer, 0.4 μmol/L RBD probe, and 10 μL 2×Premix Ex Taq Mix. The following conditions were used: 95 °C for 30 s, 40 cycles at 95 °C for 5 s, and 60 °C for 31 s in a Thermofisher QuantStudio5 and analyzed with the dedicated software.

### ELISA assay

The cell lysate, culture supernatant, and serum were homogenized for protein extraction. The spike protein RBD (Beyotime, Shanghai, China) and neutralizing antibody (Vazyme, Nanjing, China) expressions were determined using ELISA kits according to the manufacturer’s instructions.

### Measurement of RBD activity *in vitro*

We tested whether an oral vaccine for SARS-CoV-2 RBD-based DC-milk-derived exosomes (RBD-DC-milk-exos) could deliver functional RBD mRNA into human cells. For *in vitro* studies, the oral vaccine added the resulting formulations to cultures of human cells, grew the cells overnight to allow for RBD mRNA uptake and expression, and then assayed the cells for RBD activity by western blot and ELISA.

### Delivery function verification of RBD-DC-milk-exos *in vivo*

We used age-matched female BALB/c mice (Speford (Beijing) Biotechnology Co., Ltd.). All animal experimentation was performed following institutional guidelines for animal care and use and was approved by the Experimental Animal Ethics Committee of Youji (Tianjin) Pharmaceutical Technology Co., Ltd. (IACUC-20220726-05.00). Female BALB/c mice (23~26 g, Speford, Beijing, China) were housed in a pathogen-free facility with standard conditions of temperature 24 °C, a 12-hour light/dark cycle, and food and water ad libitum. Mice were employed to study the activity of RBD mRNA-DC-milk-exos administered via duodenal injection. Mice were randomly divided into 2 groups: 1) the control group (saline, 1000 μL) (N = 3); and 2) the RBD-DC-milk-exos (0.5 mg mRNA/1000 μL) group (N = 5). All treatments were started by duodenal injection on days 1, 15, and 36. All mice were sacrificed two weeks after the last treatmen with 1% isoflurane. Blood was collected at different time points (days 0, 7, 14, 21, 28, 35, 42, and 49). Verification of neutralizing antibodies of the oral vaccine for SARS-CoV-2-based milk-derived exosomes *in vivo* was executed by ELISA analysis in serum. N = 3 (control), N = 5 (RBD mRNA-DC-milk-exos), and the data are presented as mean ± STD. \*\**P* < 0.01, \*\*\**P* < 0.001 vs. control group.

### Statistical analyses

The data were represented as the mean and standard deviation. An unpaired student’s t-test was used to analyze data with only two sets. A one-way analysis of variance (ANOVA) was performed to determine whether there was a significant difference between more than two datasets, followed by Bonferroni’s post hoc test using GraphPad Prism 6.0. P < 0.05 was considered statistically significant for group differences. Asterisk (*) represented P < 0.05; a double asterisk (**) represented P < 0.01; a triple asterisk (***) represented P < 0.001.

## Acknowledgements

We thank Hangping Rui for administrative support; and also thank Fengbin Li, Mi Chen, Lin Ma, Tonglin Cui, and Xiaohan Dai for their excellent technical assistance.

## Author contributions

All authors reviewed the manuscript; X.H.G., L.L., Q.Z., M.W., and C.L.H. formulated ideas, designed the study and experiments. Q.Z., M.W., C.L.H., and Z.J.W. performed experiments and produced reagents; L.L, Q.Z., M.W., C.L.H., X.Z.M., D.L.Q., N.W., and J.H.W. analyzed experimental data; H.Q.D. supervised work. L.L. and Q.Z. wrote the manuscript. All authors contributed to the article and approved the submitted version.

## Data and materials availability

All data are available in the main text or the supplementary materials. Further inquiries can be directed to the corresponding author.

## Ethics statement

The animal study was reviewed and approved by the Experimental Animal Ethics Committee of Youji (Tianjin) Pharmaceutical Technology Co., Ltd. (IACUC-20220726-05.00).

